# Alterations of the TGFb-sequestration complex member ADAMTSL1 levels are associated with muscular defects and rhabdomyosarcoma aggressiveness

**DOI:** 10.1101/2023.03.07.531559

**Authors:** Adrien Bertrand-Chapel, Swann Meyer, Gaëtan Juban, Anita Kneppers, Paul Huchedé, Cindy Gallerne, Ruth Benayoun, Enzo Cohen, Alejandro Lopez-Gonzales, Sabrina Ben Larbi, Marion Creveaux, Lucile Vaille, Amélie Bouvier, Marine Théodore, Laura Broutier, Aurélie Dutour, Martine Cordier-Bussat, Jean-Yves Blay, Nathalie Streichenberger, Cécile Picard, Nadège Corradini, Valérie Allamand, Rémi Mounier, Perrine Castets, Marie Castets

## Abstract

Rhabdomyosarcoma (RMS) is the most frequent form of paediatric soft-tissue sarcoma and remains a medical challenge, holding in failure current therapeutic strategies. RMS shares histological features with cells of the muscle lineage and this cancer is thought to arise from malignant transformation of myogenic precursors. It has been proposed that RMS and myogenesis could represent the “Jekyll and Hyde” of skeletal muscle. The underlying idea is that some early steps of myogenic differentiation are blocked in RMS, and that understanding how the normal process has gone awry could help to decipher the biological underpinnings of tumorigenesis and tumor escape.

It is widely agreed that extracellular matrix (ECM) interferes in skeletal muscle regeneration and that defects in ECM components are clinically significant. The challenge is now to decipher actors and mechanisms responsible for the transmission of signals to muscle cells and the subsequent alterations that could be associated with RMS.

Using an original transgenic mice model, we show here that ADAMTSL1 is involved in skeletal muscle regeneration. As previously reported for other members of its family, ADAMTSL1 is part of the TGF-β-ECM-sequestering complex and likely positively regulates TGF-β-pathway activity. Last, we observed that ADAMTSL1 expression behaves as a strong prognostic factor in the aggressive fusion-positive RMS and correlates with a neural-like phenotype of tumor cells, resulting from gain of SMAD2/3/4 targets.

## INTRODUCTION

Rhabdomyosarcoma (RMS) is the most frequent form of pediatric soft-tissue sarcoma, accounting for 5% of solid pediatric tumors^1^. Two major histological subtypes are distinguished: alveolar RMS (ARMS) driven by PAX3-FOXO1 or PAX7-FOXO1 translocations in 80% of cases, and embryonal RMS (ERMS), which are genetically heterogeneous. RMS shares histological features with cells of the skeletal muscle lineage and high expression levels of myogenic proteins such as MyoD, Myogenin, and Desmin are also observed in tumors. RMS are thought to arise, at least in part, from malignant transformation of skeletal muscle precursors^2^. The notion that the molecular pathogenesis of myogenic cancers, and notably RMS, may share similarities with muscular dystrophies has begun to emerge^3^. Indeed, increased propensity to develop RMS has been reported in some murine models of dystrophies^4^. This is thought to result notably from the induction of inflammation, fibrosis and oxidative stress that accompany both tumor development and muscular dystrophies. Some of the molecules described as deregulated in dystrophies have also been connected to RMS occurrence and reciprocally. Intragenic deletions in dystrophin, mutated in Duchenne Muscular Dystrophy, are one of the underlying causes of tumor escape in RMS^5^. Proteins related to DUX4, whose gain of expression is now pointed to be largely responsible for Facio-Scapulo-Humeral Dystrophy (FSHD), are also expressed in human RMS cell lines^6,7^. Reciprocally, significant enrichment of cancer-related genes has been observed among genes altered in FSHD muscle cells, thereby suggesting that pathogenesis of dystrophies and cancers may indeed be driven by modifications of a common set of genes and associated-signaling pathways.

Among these signaling pathways, figures Transforming Growth Factor beta (TGF-β) one. TGF-β is a polypeptide that regulates several aspects of cells physiology in different tissues and has multiple effects in pathological conditions. In skeletal muscle, it has been involved in regulation of muscle growth/size, myogenesis and possibly even stemness^8^. This cytokine affects skeletal muscle function in diseases, as exemplified with fibrosis in dystrophic muscles^9^ and recently with muscle cachexia observed in some cancers^8,10,11^. TGF-β also plays a complex role in tumor progression, since it is considered as a tumor suppressor in the early steps of tumorigenesis, and as an oncogene in advanced tumors^11^. It has notably been shown to inhibit tumor cells differentiation and to promote proliferation in RMS^12^. TGF-β is secreted as a latent cytokine that is further sequestered by extracellular matrix (ECM) components. Indeed, the three TGF-β isoforms are synthetized as pro-proteins. After proteolytical cleavage, the pro-domain, called LAP (Latency-Associated Peptide (LAP)), remains non-covalently associated to the mature TGF-β moieties, forming a complex called Small Latent Complex (SLC)^13^. In the endoplasmic reticulum, this SLC complex covalently interacts with Latent-Transforming Growth Factor-β binding proteins (LTBP) to form the Large Latent Complex (LLC). Since LTBP1, 2, 3 and 4 are microfibril-associated proteins that bind to fibrillin, the LLC is incorporated to the ECM and TGF-β is stored in this latent form that is unable to bind to cell-surface receptors^13^. Once released from the LAP pro-domain, TGF-β binds to TβRI and TβRII receptors, which trigger the so-called non-canonical and canonical SMAD signaling pathway that leads to changes in gene expression.

The regulation of TGF-β bioavailability within the ECM is therefore crucial for appropriate mobilization and activation of this cytokine. Deciphering the mechanisms underlying the mobilization of latent TGF-β from this local reservoir could shed new light on the role exerted by ECM on skeletal muscle homeostasis on one hand, and on alterations associated to tumorigenesis in RMS on the other hand. Very little is known about alterations of ECM in RMS, but some ECM proteins have been characterized as biomarkers^14^.

Here, we decided to study the role of ADAMTSL1 [a disintegrin and metalloprotease domain with thrombospondin motifs)-like 1] in these processes. ADAMTSL1 is the first identified member of a family of ECM-associated proteins that are closely related to ADAMs and ADAMTS metalloproteinases, but devoid of enzymatic activity^15^. The role of ADAMTSL1 in mammals remains unknown. Interestingly, high expression levels of ADAMTSL1 were detected/described in skeletal muscle^15^. Moreover, 3 of the 7 members of the ADAMTSL family, namely ADAMTSL2, 3 and 6, are part of the ECM complex regulating TGF-β bioavailability^16–18^. In line with these issues, we show here that ADAMTSL1 contributes to skeletal muscle homeostasis by modulating TGF-β bioavailability, and that the subsequent impairment of its function is involved in RMS phenotype, and notably behaves as a prognostic factor in the most aggressive fusion-positive ones.

## RESULTS

### Loss of ADAMTSL1 induces mild spontaneous muscular phenotype during ageing but is involved in muscle regeneration as a FAP-secreted protein

Since ADAMTSL1 was described as mostly expressed in skeletal muscles^15^, we analysed its function in this tissue by characterizing an original *Adamtsl1* KO mice, generated *via* MMRRC repository, through constitutive excision of exon 1 (Figure 1A). We validated the deletion of *Adamtsl1* exon 1 in progeny using specific primers (Figure S1A). Adamtsl1^+/-^ and Adamtsl1^-/-^ mice were born at expected mendelian frequencies, healthy and fertile.

**FIGURE 1:**
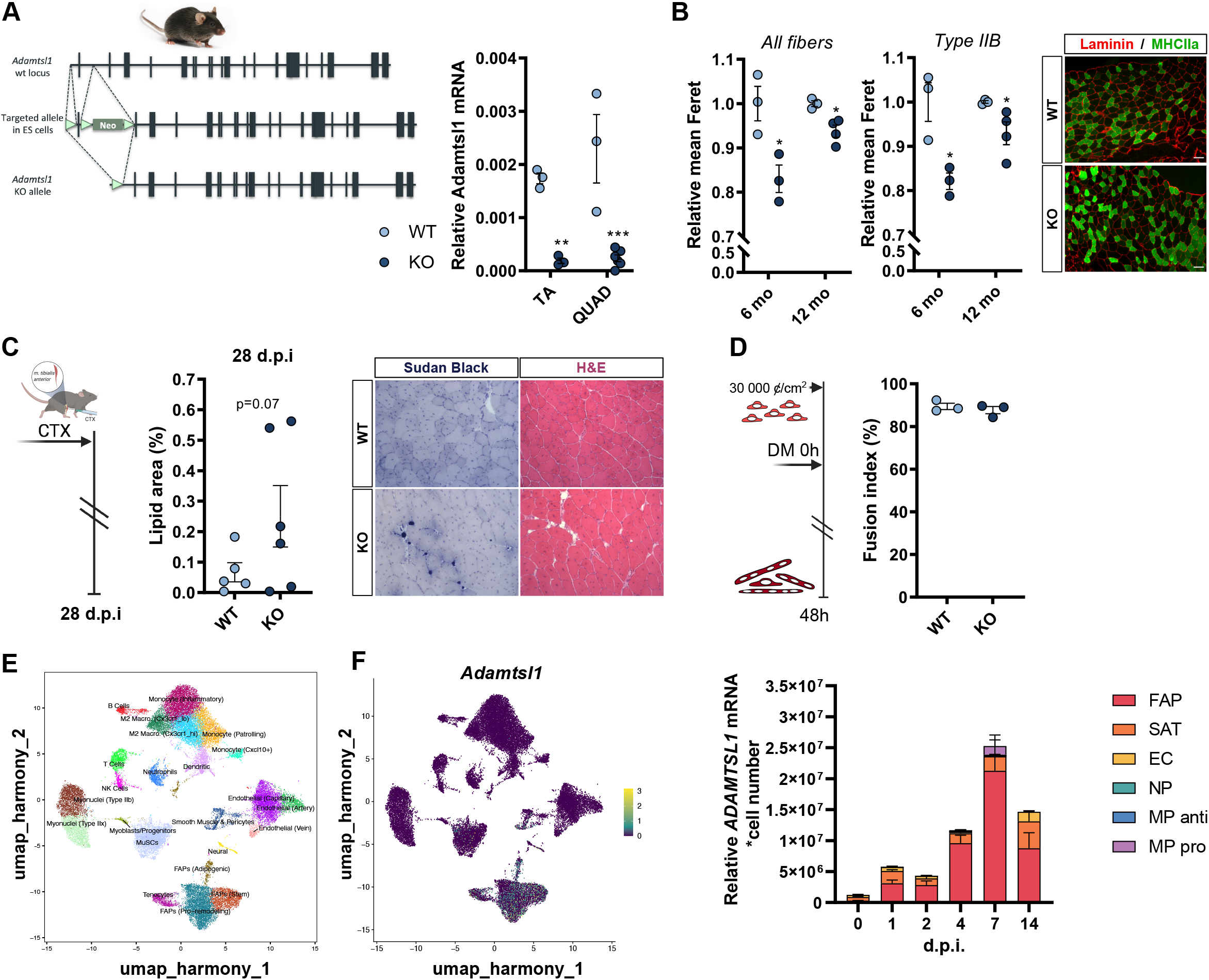
Loss of ADAMTSL1 induces mild skeletal muscle regeneration defects in mice. **A**. Scheme of *Adamtsl1* exon 1 KO strategy and impact on expression of its expression in KO versus WT mice. RT-QPCR were performed on mice Quadriceps (QUAD) and Tibialis Anterior (TA) muscles. Results are mean +/-std, n=3. **: p<0.01, ***: p<0.001, t-test. **B**. Mean fiber size showing slight hypotrophy in ADAMTSL1 KO mice muscle at 6 and 12 months of age (left panels). 6mo: 6months-old mice. 12 mo: 12 months-old mice. Results are mean+/-std, indexed to mean of control animals; n=3, *: p<0.05, t-test. Representative images of Laminin/MYH4 stainings are shown (right panel). **C**. Impaired muscle regeneration in *Tibialis Anterior* muscle of *Adamtsl1* KO mice after an injury induced by cardiotoxin injection. Quantification of lipid droplets infiltrates in KO mice (n=4) compared to controls (n=4) (left panel). Results are means +/-std; p=0.07, t-test. Representative images of muscle histology and lipid droplets infiltrates (Sudan Black staining) in KO mice compared to controls are shown (right panel). D.p.i: Days post injury, H&E: Hematoxylin & Eosin, CTX: cardiotoxin. **D**. Absence of significant changes in the fusion index of satellite cells derived from ADAMTSL1 WT and KO mice. DM: Differentiation medium. Results are means +/-std. **E**. UMAP visualization of unified scRNA-seq data of skeletal muscle cell populations (left panel), showing that ADAMTSL1 is mostly expressed in Fibro-Adipogenic Precursors (FAP) (right panel). **F**. RT-qPCR analysis of ADAMTSL1 mRNA level in various cell populations FACS-sorted from *Tibialis Anterior* muscle. Muscles were analysed 0, 1, 2, 4, 7 and 14 days after injury (d.p.i).

We confirmed that the transcription of *Adamtsl1* is significantly turned off in skeletal muscles of homozygous KO animals as compared to their wild-type (WT) littermates (Figure 1A). Of note, ADAMTSL1 KO mice are identifiable by a kink at the end of the tail (Figure S1B). Loss of ADAMTSL1 is not associated with obvious changes in skeletal muscle organization in young animals (data not shown). However, defects can be observed in KO animals from 6 months of age, with a 20% decrease in mean muscle fiber diameter, especially in fast contraction type IIb fibers (Figure 1B). This muscle fiber hypotrophy is associated with a slight atrophy of the *Gastrocnemius*+*Soleus* assembly at 12 months only (Figure S1C). Consistent with progressive muscle weakening during ageing, a typical hunchback is observable in one-year Adamtsl1^-/-^ animals (Figure S1D). These changes were not associated with any difference in the number of Pax7^+^ cells nor in the percentage of centronucleated fibers, suggesting that the defects may likely result in part from alterations of muscle cell differentiation (Figure S1E-F). We then wondered what its contribution in muscle regeneration could be. To do so, we compared skeletal muscle regeneration after cardiotoxin (CTX) injection in adult KO mice and their wild-type counterparts. As shown on Figure 1C, ADAMTSL1 loss is associated with defects in muscle regeneration characterized by lipid droplets accumulation. These defects are not due to intrinsic failure of *Adamtsl1* KO satellite cells to differentiate and fuse, as shown by their ability to do so when isolated *in vitro* (Figure 1D). Of note, the number of Pax7^+^, a readout of satellite cell self-renewal, are similar in KO and WT animals 28 days post-CTX injection (Figure S1G). Since satellite cells phenotype does not seem to be intrinsically affected by ADAMTSL1 loss, we sought to identify whether this matrix protein was secreted by a specific skeletal muscle cell population. Using single-cell analysis, we observed that ADAMTSL1 is mostly expressed by Fibro-Adipogenic Precursors (FAP, Figure 1E) and that its expression is increased during regeneration in this population (Figure 1F).

Then, loss of FAP-produced ADAMTSL1 is associated with muscular defects during ageing and regeneration.

### ADAMTSL1 belongs to the FBN1 matrix complex regulating TGF availability

To clarify the role of ADAMTSL1 in muscle homeostasis, we decided to study it in the context of muscular dystrophy. Indeed, repeated lesions of muscle fibers in patients suffering from Duchenne dystrophy trigger permanent regeneration cycles, finally leading to muscle fibrosis^19^. By integrating public transcriptomic data from the analysis of muscles’ biopsies of Duchenne patients at different stages of the disease, we observed that ADAMTSL1 expression increases sequentially at the three stages of the disease, *i*.*e*. mild, moderate and severe (Figure 2A). The same trend is observed for ADAMTSL3 and 4, although less significantly, but not for ADAMTSL2 (Figure S2A). On the contrary, the expression of ADAMTSL5 decreases under the same conditions. Interestingly, ADAMTSL2, 3 and 6 have been described as matrix proteins, associated with the TGF-β-fibrillin sequestering complex^16–18^, and TGF-β is one of the main contributors to the muscle fibrosis observed in the context of muscular dystrophy^20^. Thus, it was tempting to speculate that ADAMTSL1 could play a role in muscle homeostasis by regulating TGF-β bioavailability. As shown on Figure 2B, ADAMTSL1 colocalizes with LTBP1 and FBN1 in the muscle ECM of patients with dystrophy. In addition, we used the PROGENy method to infer downstream apoptotic-response footprint from perturbation-response genes indicative of the TGF-β activity^21^. This score is significantly increased in moderate grade dystrophic muscle samples as compared to mild grade, and even more significantly between severe and mild grades (Figure S2B), consistent with the activation of this pathway in this pathological context^22^. As shown in Figure 2C, TGF-β pathway activity is significantly correlated with ADAMTSL1 expression notably in the group of patients with mild dystrophy (Figure 2C). These data suggest that ADAMTSL1 may play a positive role in the regulation of TGF-β bioavailability in muscles and are consistent with analyses performed in mice showing that the expression of 18% of the major target genes of the TGF pathway is significantly altered in muscles from KO mice as compared to wild-type mice (Figure 2D).

**FIGURE 2:**
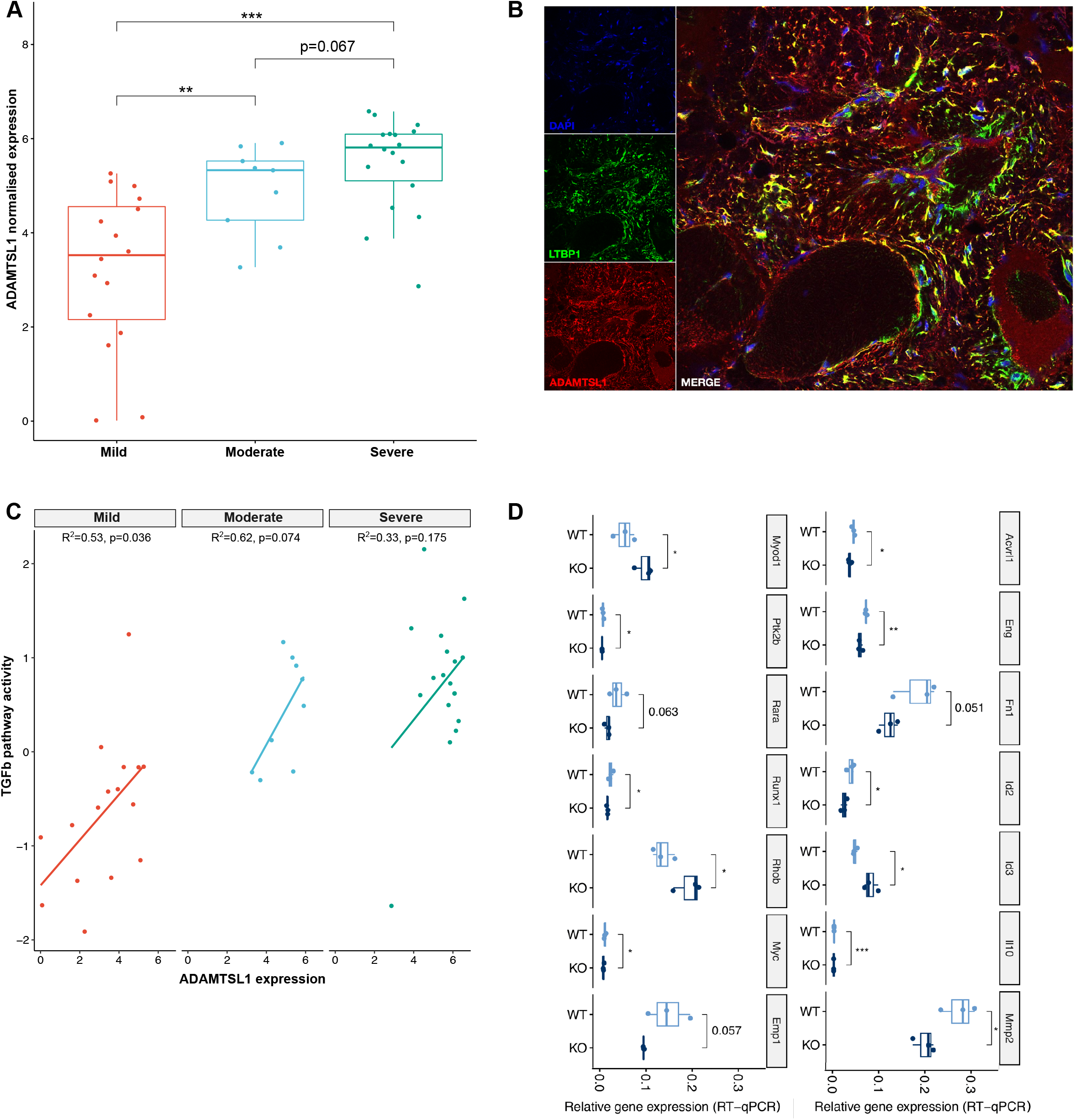
ADAMTSL1 is linked to the regulation of TGF-β pathway in skeletal muscles. **A**. Expression of ADAMTSL1 in skeletal muscle samples from patients with Duchenne dystrophy. Analysis was made from GSE109178 microarray cohort (cohort 1). Disease grade was reported in clinical data. **: p<0.01, ***: p<0.001, t-test. **B**. Colocalization of ADAMTSL1 and LTBP1 in skeletal muscles from patients with muscle dystrophy. ADAMTSL1 and LTBP1 are respectively stained in red and green. Nuclei are counterstained in blue. Representative confocal images of a muscle biopsy from a patient with Ullrich Congenital Muscular Dystrophy are shown. **C**. Correlation between ADAMTSL1 expression and TGF-β pathway activity inferred from a specific gene response signature using PROGENy algorithm in skeletal muscles from patients with Duchenne dystrophy (cohort 1). The Pearson correlation coefficient is shown for each disease grade, with its associated p-value. **D**. Modifications of TGF-β target genes’ expression in ADAMTSL1 KO versus WT animals. TGF-β targets were quantified by Q-RT-PCR in skeletal Tibialis Anterior muscle from 3 months-old mice, using the dedicated kit from Qiagen. Results are mean+/-std; n=3. *: p<0.05, one-sided t-test.

Altogether, these data indicate that ADAMTSL1 likely replays a role in muscle regeneration *via* regulation of TGF-β bioavailability.

### ADAMTSL1 expression positively correlates with prognosis in fusion-positive rhabdomyosarcoma and with expression of SMAD3/4 neural-like targets

RMS are thought to result at least in part from defects in the process of myogenesis, with a premature arrest of muscle cell differentiation^1,23,24^. Considering the role of TGF-β in tumorigenesis and its impact on muscle differentiation in RMS^12,25,26^, we then wondered whether alterations in ADAMTSL1 expression might play a role in RMS. Integration of RNAseq data indicates that ADAMTSL1 expression is significantly lower in RMS than in skeletal muscle samples, in two independent cohorts (Figure 3A and Figure S3A). To go further, we analyzed the clinical value of ADAMTSL1 expression. Interestingly, the integration of transcriptomic data indicates that ADAMTSL1 expression is positively correlated with the prognosis in the so-called fusion-positive RMS (FP-RMS), expressing the Pax3/7-FOXO1 translocation, but not in the fusion-negative RMS (FN-RMS), which lack it (Figure 3B-C). Similarly, a high level of methylation of the *ADAMTSL1* locus is associated with a poor prognosis in FP-RMS, but not in FN-RMS (Figure S3B-C). To clarify the molecular identity and origin of the prognostic differences observed between tumors with high and low levels of ADAMTSL1, we performed a differential expression (DE) analysis between these two subgroups. Of the 270 DE genes corresponding to the intersection of two independent cohorts (Figure S3D), we observed that 64.7% are more expressed in the ADAMTSL1-high tumors group as compared to the ADAMTSL1-low group, as shown on heatmaps (Figures 3D and S3E). Functional gene set enrichment indicate that the ADAMTSL1-high tumors group is robustly defined by neural-related pathways, especially some involved in regulation of cell migration/adhesion (Figure 3E). Last, RMS tumors expressing high level of ADAMTSL, but not those with low levels of this gene, are significantly enriched in target genes of SMAD2/3/4, which are the transcription factors activated downstream of TGF-β (Figure 3F).

**FIGURE 3:**
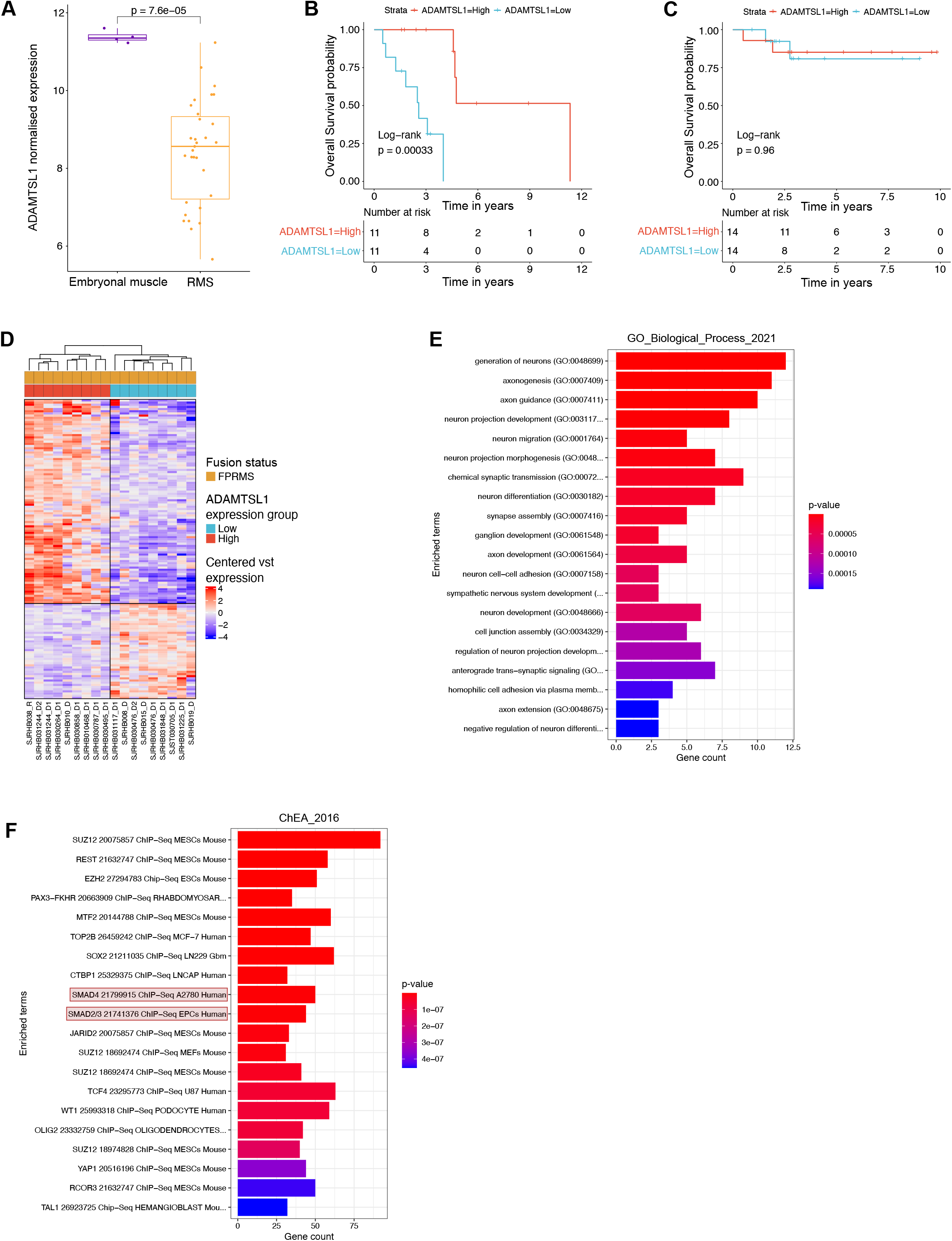
ADAMTSL1 expression is a good prognostic marker associated with neural-like phenotype in FP-RMS. **A**. Expression levels of ADAMTSL1 between normal muscles and FP-RMS samples (cohort 2, RNAseq). Differences between groups were tested using wilcoxon signed-rank test with associated statistical probability displayed on top. **B-C**. Survival analyses of ADAMTSL1 expression value in FP-RMS samples (B) and FN-RMS samples (C) (cohort 4). Kaplan-Meier curves were generated between ADAMTSL1-high and -low groups, generated as defined in Material and Methods section. Differences of overall survival probability between both groups were tested using log-rank tests and associated statistical probabilities are displayed on the graph, for both FP- and FN-RMS cohorts. Number of patients at risk are indicated in the tables below the curves. **D**. Heatmap representing the transcriptomic expression levels of apoptotic genes significantly differentially expressed between ADAMTSL1-high and low groups of FP-RMS (cohort 4). Normalized and scaled gene expression levels are color-coded with a blue (low expression) to red (high expression) gradient. Samples in columns are clustered using Ward’s method on the inverse Spearman’s correlation coefficient matrix. **E**. Functional enrichment between ADAMTSL1-high and -low groups of FP-RMS (see Material and Methods). GO_Biological_Process_2021 Enrichment reveals a significant enrichment in neural-related pathways in ADAMTSL1-high group. **F**. Functional enrichment between ADAMTSL1-high and -low groups of FP-RMS (see Material and Methods). ChEA_2016 Enrichment reveals a significant enrichment notably in SMAD2/3/4 targets in ADAMTSL1-high group.

Then, ADAMTSL1 behaves as a prognostic factor in FP-RMS and likely interfere with tumor cell properties through regulation of TGF-β pathway activity.

## DISCUSSION

RMS account for 5% of all pediatric tumors. Intensive therapeutic management has improved their 5-year survival from 25% in 1970 to 60-70% since 2000^2^. However, outcome has reached a plateau since that time, despite many randomized chemotherapy trials. Moreover, 2/3 of cured children suffer from long-term side effects of both disease and treatments^2,27^. This underscores the need to elucidate molecular mechanisms of these malignancies to develop new therapeutic strategies, both beneficial for patients in therapeutic failure and with reduced toxicity.

It has been proposed that RMS and myogenesis could represent the “Jekyll and Hyde” of skeletal muscle^1^. The underlying idea is that some early steps of myogenic differentiation are blocked in RMS: understanding how the normal process has gone awry could help to decipher the biological underpinnings of tumorigenesis and of tumor escape.

Here, we show that ADAMTSL1, whose function was yet undefined, likely plays a role in muscle regeneration as a matrix protein, through regulation of the TGF-β pathway, and that alteration of its expression has a direct impact on FP-RMS aggressiveness.

The role of ECM in skeletal muscle regeneration is not well understood, but it is widely agreed that alterations in ECM components are clinically significant. Along this line, two other members of the ADAMTSL family, namely ADAMTSL2 and 6, have been linked to genetic diseases of the Marfanoid spectrum, with mutations identified respectively in geleophysic dysplasia and thoracic aortic aneurysm^16,28^. Although not related to skeletal muscular defects, both proteins were also shown to be part of the FBN1 complex regulating the bioavailability of TGF-β^17^. Moreover, ADAMTSL2 was directly identified as a negative regulator of myofibroblast differentiation through inhibition of TGF-β^29^. Similarly, KO of ADAMTSL3 is associated with severe cardiac defects in mice, which result from an increase in TGF-β signalling, subsequently impairing myofibroblasts differentiation upon heart failure^18^. While these data suggest a conserved role of the 6 members of the ADAMTSL family in the regulation of TGF-β release, our results indicate that ADAMTSL1 may play the opposite role, by acting as a positive regulator of the TGF-β signalling pathway: ADAMTSL1 could thus promote TGF-β release rather than its sequestration in the matrix. Consistently, fatty deposits similar to those observed in ADAMTSL1 KO mice after injury were observed in transgenic mice that overexpress the human TGF-β-sequestering LTBP4 protein^30^, suggesting that ADAMTSL1 and LTBP4 may have antagonistic function on TGF-β pathway. Moreover, ADAMTSL1 expression correlates with TGF-β activation level in patients with Duchenne dystrophy, and its expression increases with disease severity: it remains however to be determined whether gain of ADAMTSL1 is causally involved in muscle degeneration observed in this pathology, or if it is a consequence of muscle fibrosis.

Presence of a regenerative microenvironment in skeletal muscle, coupled with oncogenic alterations in cells, was shown to be sufficient to favor FN-RMS tumorigenesis in mice^31^. FBN2 was also shown to have a prognostic value in FN-RMS, although through a yet undefined mechanism, but further highlighting the clinical relevance of ECM defects in RMS^14^. In contrast to the gain of ADAMTSL1 observed in dystrophic muscle samples, we show that the loss of ADAMTSL1 is highly significantly correlated with tumor aggressiveness but solely in FP-RMS, which are the most resistant to treatments^2^. Of note, this prognostic value is not only visible at the transcriptomic level, but also at the DNA methylation one, suggesting that the alteration of ADAMTSL1 expression in tumors may result from epigenetic modifications. Bioinformatic analyses indicate that ADAMTSL1 expression is positively correlated with a neural-like phenotype, which has been previously described by others in FP-RMS^24^. It had been established that the neuronal guidance activity of MADD-4, the C. elegans ortholog of ADAMTSL1, was dependent on the presence of the netrin-1 receptor, DCC^32^. Interestingly, DCC is one of the neuronal genes significantly enriched in the ADAMTSL1-high tumor group, suggesting that some mechanisms of action may be conserved across species. The oncogenic valence of the FP-RMS neural state has not been described so far. Further analyses are however needed to understand how this observation may be compatible with the better prognosis of FP-RMS with the highest ADAMTSL1 expression. Along this line, it was previously shown that ADAMTSL1 may play an oncogenic role in chondrosarcoma through positive regulation of proliferation^33^, suggesting a pleiotropic role of this protein on tumor cell phenotype, which is probably an indirect consequence of the complex role of TGF-β in tumor initiation and escape^11^.

If the precise role of ADAMTSL1 in muscular dystrophies and RMS remains to be fully established, our work makes it a protein worth studying to understand how deregulations of the skeletal muscle matrix composition can create a deleterious microenvironment.

## MATERIAL AND METHODS

### Design of an ADAMTSL1 knock-out model

The B6;129S5-Adamtsl1tm1Lex/Mmucd mouse strain presenting a constitutive deletion of the first exon of ADAMTSL1 was obtained from MMRRC. In brief, targeting vector was constructed using a fragment of ADAMTSL1 gene around exon 1 (NCBI accession: AK045085). Embryonic stem cell electroporation, selection, and culture, as well as generation of chimaeric mice and Southern blot analysis were performed in the MMRRC according to classical procedures (https://www.mmrrc.org/catalog/sds.php?mmrrc_id=32134). Heterozygous mice were then backcrossed at least 10 times in a C57BL/6 background. Double heterozygous ADAMTSL1^+/-^ mice of the offspring were interbred to generate mice homozygous for the ADAMTSL1 exon 1 deletion. Routine genotype analysis of mice was performed by PCR assay on DNA purified from tail biopsies (Extract N-Amp Tissue PCR kit, Sigma Aldrich). Wild-type and KO mutant alleles were distinguished using primers available on request. CT-Scan was performed on the animal imaging platform of CRCL. All experiments were performed in accordance with the relevant guidelines and regulations of the animal ethics committee (Authorization no. CLB-2015-024; accreditation of laboratory animal care by CECCAPP, ENS Lyon-PBES).

### Quantitative RT–PCR

Total RNA was extracted with the RNeasy Fibrous Tissue Mini Kit (Qiagen) and 1 μg was reverse transcribed using the iScript cDNA Synthesis kit (Bio-Rad). Real-time quantitative RT– PCR was performed on a LightCycler 2.0 apparatus (Roche) using the Light Cycler FastStart DNA Master SYBERGreen I kit (Roche). Oligonucleotide sequences are available on request. Profiling of TGF-β targets was performed using the RT^2^ Profiler™ PCR Array Pig TGF-β Signaling Targets kit from Qiagen following manufacturers’ instructions.

### Skeletal muscle injury

Prior injection of CTX (12 mM in saline, Latoxan#L8102), mice were anesthetized by intraperitoneal injection of ketamine at 0.1 mg/g body weight and xylazine at 0.01 mg/g body weight diluted in saline solution. 50 μl of CTX was then injected into *Tibialis Anterior* muscles to induce injury, and mice were euthanized 0, or 28 days after the injury.

### Satellite cell cultures – *in vitro* model of adult myogenesis

Skeletal muscle myoblasts derived from WT and KO ADAMTSL1 mice were isolated using the Satellite Cell Isolation Kit MACS protocol (Miltenyi Biotec), as described previously^34^.

MACS isolated Satellite cells were seeded onto Gelatin (Sigma #G1393) coated supports at 3000 cells/cm^2^ and amplified ≤3 passages in proliferation medium. After amplification, MuSC progeny (*i*.*e*., myoblasts) were trypsinized and plated at indicated densities onto Matrigel coated supports. To induce differentiation, proliferation medium was removed and replaced by differentiation medium (DMEM F12, 2% Horse serum (HS) (Gibco #16050130), and 1 % PS).

### Histological and immunohistochemistry analyses on muscle cryosections and cell cultures

For immunohistochemical analyses, muscles of interest were isolated, embedded in tragacanth gum (VWR #ICNA0210479280), frozen in liquid nitrogen cooled isopentane, and stored at -80°C until use. 10 μm-thick cryosections were prepared for hematoxylin-eosin (HE) staining and immunolabeling. HE staining was used to assess the efficiency of CTX injections, and muscles were only kept for further analyses if ≥75% of the muscle area consisted of centrally nucleated fibers.

For lipid coloration, sections were fixed in 4% formaldehyde at 4°C during 30 min, and then stained with Sudan Black (Sigma-Aldrich Corp, St. Louis, MO, USA) for 60 minutes. Light microscopy was performed using an upright microscope (Leica and Olympus), and pictures were captured using a monochrome camera (DS-Ri1, Nikon).

For immunohistochemistry, muscle cryosections were stained with ADAMTSL1 (Invitrogen MA1-41249), LTBP1 (Abcam ab78294), FBN1 (Abcam ab53076), Laminin (Abcam ab11575) and MHCIIA (DSHB SC-71) antibodies as described previously^36^. They were mounted in Vectashield DAPI (Vectorlabs) and observed with an Axiophot microscope (Zeiss). Images were captured using the MetaView software (Ropper Scientific). For identification of satellite cells, sections were fixed for 20 min in 4% paraformaldehyde (PFA) at Room Temperature (RT), and permeabilized for 6 min in 100% methanol at -20°C. After antigen retrieval for 2*5 min in 10 mM Citrate buffer at 90°C, sections were saturated in 4% BSA for 2 hr at RT. Sections were then co-labelled overnight at 4°C with primary antibodies directed against Paired Box 7 (Pax7; DSHB; 1:50) and Laminin α1 (Sigma-Aldrich #L9393; 1:100) in 2% BSA. Secondary antibodies were coupled to FITC, Cy3 or Cy5 (Jackson ImmunoResearch Inc.; 1:200) and incubated for 45-60 min at 37°C. Nuclear counterstain was performed by 10 sec incubation with Hoechst (Sigma-Aldrich #14533; 2 μM), and coverslips were mounted with Fluoromount G (Interchim, #FP-483331).

For fusion assay, cultured cells were fixed for 10 min in 4% PFA at RT, permeabilized for 10 min in 0.5% Triton X-100 in PBS, and saturated in 4% BSA for 1 hr at room temperature (RT). Cells were then incubated overnight at 4°C with primary antibodies directed Desmin (Abcam #Ab32362; 1:200) in 2% BSA. Secondary antibodies were coupled to FITC, Cy3 or Cy5 (Jackson ImmunoResearch Inc.; 1:200) and incubated for 45-60 min at 37°C. Nuclear counterstain was performed by 10 sec incubation with Hoechst (Sigma-Aldrich #14533; 2 μM), and cells were covered with Fluoromount G (Interchim, #FP-483331).

Randomly chosen fields of view from sections or cells cultured on removable chamber slides were acquired using a Zeiss Axio Imager Z1 connected to a Coolsnap Myo camera. For each condition of each experiment, at least 5-10 randomly chosen fields of view were counted. The fusion index was calculated as the percentage of nuclei within a cell containing ≥2 nuclei.

### RT-qPCR analysis in various cell populations FACS-sorted from *Tibialis Anterior* muscle

Muscle cell populations were FACS-sorted as previously described^37^. Briefly, TA muscles were dissociated and digested in DMEM/F12 medium (GIBCO) containing 10 mg/ml of collagenase B and 2.4 U/ml Dispase II (Roche Diagnostics) for 30 min at 37°C under agitation and passed through a 30 μm cell strainer (Myltenyi Biotec). CD45^pos^ and CD45^neg^ cells were separated after incubation with mouse CD45 MicroBeads (Myltenyi Biotec) using a MS column (Myltenyi Biotec) on a MiniMACS Separator (Myltenyi Biotec). CD45^pos^ cells were incubated with mouse FcR Blocking Reagent (Myltenyi Biotec) for 20 min at 4°C in PBS 2% FBS and stained with PE-Cy7-conjugated anti-CD45 (25-0451-82, eBioscience), Alexa Fluor 647-conjugated anti-CD64 (139322, Biolegend) and PE-conjugated anti-Ly6C (12-5932-82, eBioscience) antibodies for 30 min at 4°C. CD45^neg^ cells were stained with PE-Cy7-conjugated anti-CD45, PerCP-Cy5.5-conjugated anti-Sca-1 (45-5981-82, eBioscience), Alexa Fluor 647-conjugated anti-α7-integrin (AB0000538, AB lab, University British Columbia), PE-conjugated anti-CD31 (12-0311-82, eBioscience) and FITC-conjugated anti-CD34 (11-0341-82, eBioscience) antibodies for 30 min at 4°C. After addition of DAPI (BD Biosciences) to exclude dead cells, live cells were sorted using a FACS Aria II cell sorter (BD Biosciences).

Total RNA was isolated using NucleoSpin RNA Plus XS kit (Macheray-Nagel) and retro-transcribed into cDNA with SuperScript II Reverse Transcriptase (Invitrogen). Quantitative PCR was realized in triplicate on a CFX Connect Real-Time PCR Detection System (Bio-Rad) using LightCycler 480 SYBR Green I Master kit (Roche Diagnostics). Calculation of relative expression was determined by the Bio-Rad CFX Manager™ software and fold change was normalised as Normalised Relative Quantity (or ΔΔCq) for each series, as 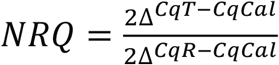 where T is the target sample, Cal is the calibrator value (*i*.*e*. the mean of all sample Cqs of the series) and R is the housekeeping gene Cyclophilin A. Sequence of primers used were as follows: 5’-TGGCTCAGAATGAAGCAGGG-3’ (Adamtsl1 fwd), 5’-AGTAAACACCTGAGTCGCGG-3’ (Adamtsl1 rev), 5’-GTGACTTTACACGCCATAATG -3’ (Cyclophilin A fwd) and 5’-ACAAGATGCCAGGACCTGTAT -3’ (Cyclophilin A rev).

### Cell culture

3T3 and RH30 were cultured according to manufacturers’ instructions. 3T3 stably expressing the short isoform of ADAMTSL1 tagged with GFP (available on request) were established using JetPrime and selected on the basis of their resistance to puromycin.

### Public scRNA-seq data of muscle tissue

UMAP of cell types and ADAMTSL1 expression were made using harmony integrated data collected by the Single-Cell Muscle Project (scMuscle)^38^, which is available for download at https://datadryad.org/stash/dataset/doi:10.5061%2Fdryad.t4b8gtj34.

### Public microarray (gene expression and methylation) datasets

GSE109178 (cohort 1, DMD samples) and GSE28511 (cohort 3, skeletal muscle and RMS samples) datasets were downloaded on GEO (www.ncbi.nlm.nih.gov/geo/), then log2-transformed. Raw CEL files of ETABM-1202 dataset^39^ (cohort 4) were downloaded on ArrayExpress (https://www.ebi.ac.uk/biostudies/arrayexpress/studies/E-TABM-1202), and were normalized using the Robust Multiarray Average (RMA) algorithm (oligo R library v 1.58.0)^40^. The GSE167059 methylation dataset (cohort 5) used for survival analysis was downloaded on R2 Cancer (https://hgserver1.amc.nl/cgi-bin/r2/main.cgi).

### Generation and processing of RNA-seq datasets

#### St Jude RMS

HTseq count files of St Jude’s RMS tumors (cohort 6) were downloaded on the St Jude Cloud platform (https://www.stjude.cloud) and generated as described^41^.

#### Inhouse RMS cohort (cohort 2)

Tissue banking and research were conducted according to national ethics guidelines, after obtaining the written informed consent of patients. For RNA-seq library construction of cohort 3, total RNAs from tissues were isolated using the AllPrep® DNA/RNA FFPE kit (Qiagen, cat. no. 80224) following manufacturer’s instructions. Libraries were prepared with Illumina Stranded mRNA Prep (Illumina, cat. no. 20040534) following recommendations. Quality was further assessed using the TapeStation 4200 automated electrophoresis system (Agilent) with High Sensitivity D1000 ScreenTape (Agilent). All libraries were sequenced (2×100 bp) using Agilent SureSelect RNA XTHS2 All Exon V8 (Agilent, cat. no. G9991A) according to the standard Agilent protocol.

Quality control of reads was performed using FastQC (v.0.11.9)^42^, followed by trimming of Illumina adapter sequences with Cutadapt (v.3.4)^43^ using the -a CTGTCTCTTATACACATCT and -A CTGTCTCTTATACACATCT parameters. Reads were mapped to the GRCh38 human genome using “two-pass” mode STAR (v.2.7.9)^44^, with Ensembl v104 annotations. Gene counts were then computed using HTseq-count (v.0.13.5)^45^ with the following parameters: “--order pos” and “--stranded reverse”.

#### Data normalization of both datasets

HTseq count files were then loaded in R (v 4.2.0) and two filtration steps were applied using the annotations of org.Hs.eg.db (v 3.15.0)^46^: genes with low counts (less than 10 reads across samples) were removed; non-protein-coding genes were removed. Filtered gene counts were converted into a DESeq2^47^ object with the design parameter set to account for the sample type (RMS/embryonal muscle/fetal muscle) in the case of the Inhouse cohort, and for the fusion status (FNRMS/FPRMS) in the cohort 6. Counts were then normalized with vst using “blind = FALSE”.

To remove unwanted variability driven by technical factors in the cohort 6, the removeBatchEffect function (limma R library v 3.50.3)^48^ was used with the original dataset specified as the variable to consider in the linear model.

The package ggplot2 (v 3.3.5)^49^ was used for graphical representation.

### Comparison of ADAMTSL1-High and ADAMTSL1-Low samples

#### Definition of ADAMTSL1 expression groups

In both cohorts 4 and 6, patients were stratified according to ADAMTSL1 expression level: samples with ADAMTSL1 levels > 3^rd^ quartile (Q3) formed the high expression group, while samples with ADAMTSL1 levels < 1^st^ quartile (Q1) formed the low expression group. For cohort 5, the KaplanScan method implemented by R2 was used to identify the cutoff of the ADAMTSL1 probe cg14523394, separately for each histology (eRMS/aRMS).

#### Survival analysis

Kaplan-Meier survival analysis was run on cohorts 4 and 5 (ETABM-1202 and GSE167059) for which clinical data were available, using the R packages survminer (v 0.4.9)^50^ and survival (v 3.3-1)^51^. Log-rank test was used to assess significance between the survival probability of ADAMTSL1 High samples and ADAMTSL1 Low samples, in FNRMS and FPRMS separately, for all survival analysis.

#### TGF-β pathway activity

The R package PROGENy (v 1.18.0)^21^ was used to assess the activity of the TGF-β pathway using default parameters. Boxplots were made with ggplot2 (v 3.3.5)^49^.

#### Differential expression analysis and enrichment

Differential expression analysis on cohort 4 was run with the R package limma (v 3.50.3) based on previously defined ADAMTSL1 expression groups, separately for FNRMS/FPRMS. Genes a with a raw p-value cutoff of 0.05 were considered as differentially expressed (DE), yielding 1095 DE genes. For RMS cohort 6, DESeq2 was used to identify differentially expressed genes between ADAMTSL1-high and ADAMTSL1-low samples, separately in FNRMS and FPRMS. The alpha parameter of the “results” function was set to 0.05, and DE genes were identified with an adjusted p-value (FDR) cutoff of 0.05.

The EnrichR package (v 3.0)^52^ was used separately on DE genes upregulated in ADAMTSL1-high samples and DE genes upregulated in ADAMTSL1-low samples, with the following databases: Gene Ontology (GO) Biological Process 2021, GO Molecular Function 2021, GO Cellular Compartment 2021, KEGG 2021 Human, MSigDB Hallmarks 2020, Reactome 2016 and ChEA 2016.

#### Correlation analysis

Correlations between ADAMTSL1 and all other genes in each fusion subtype (FPRMS/FNRMS) were assessed using Pearson’s coefficient. Genes were considered significantly correlated when unadjusted p-value < 0.05. Venn diagram representation of the significantly correlated genes identified was made with ggvenn (v 0.1.9)^53^.

## Statistical analyses

For *in vitro* experiments, replicates signify independent experiments. For *in vivo* experiments, replicates signify individual muscles. All results were analysed using unpaired parametric analyses. Statistical significance was determined using two-sided Student’t *t* tests, or Welch *t* test in case of unequal variance.

## Data availability

GSE109178 (cohort 1) and GSE28511 (cohort 3) datasets were downloaded on GEO (www.ncbi.nlm.nih.gov/geo/). Cohort 5 data was downloaded on R2 Cancer (https://hgserver1.amc.nl/cgi-bin/r2/main.cgi). Raw CEL files of cohort 4 (accession code ETABM-1202) were downloaded on ArrayExpress (https://www.ebi.ac.uk/biostudies/arrayexpress). St Jude dataset (cohort 6) was downloaded on the St Jude Cloud platform (https://www.stjude.cloud). Newly generated RNA-seq data of the Inhouse cohort will be made available on GEO (cohort 2).

## Acknowledgements

We thank the patients and their families who consented to participate in this study, as well as the Ligue Nationale contre le Cancer, and the Ligue Contre le Cancer du Rhône, which provided the funding for this project. We also thank clinical teams from IHOPe (Institut d’Hématologie et d’Oncologie Pédiatrique) and HFME (Hôpital Femme Mère Enfant – HCL) for their support and contributions, as well as all the facilities from the CLB and CRCL. We are grateful to Séverine Tabone-Eglinger, Loïc Sebileau and Anne-Sophie Bonne for their help with the management of regulatory procedures.

