## Supplementary figures and images for "Alterations of the TGFb-sequestration complex member ADAMTSL1 levels are associated with muscular defects and rhabdomyosarcoma aggressiveness"

### Supplemental Figures

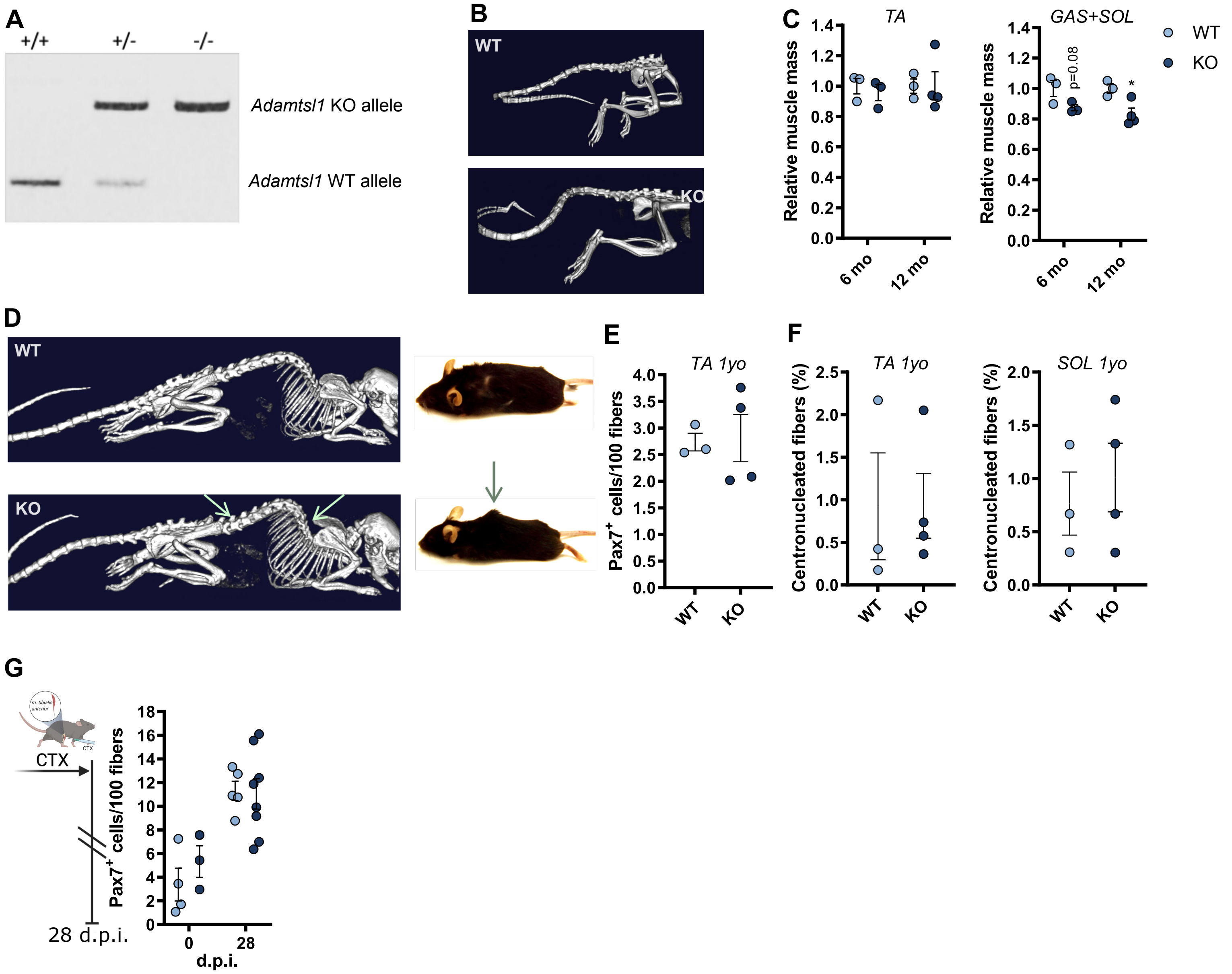

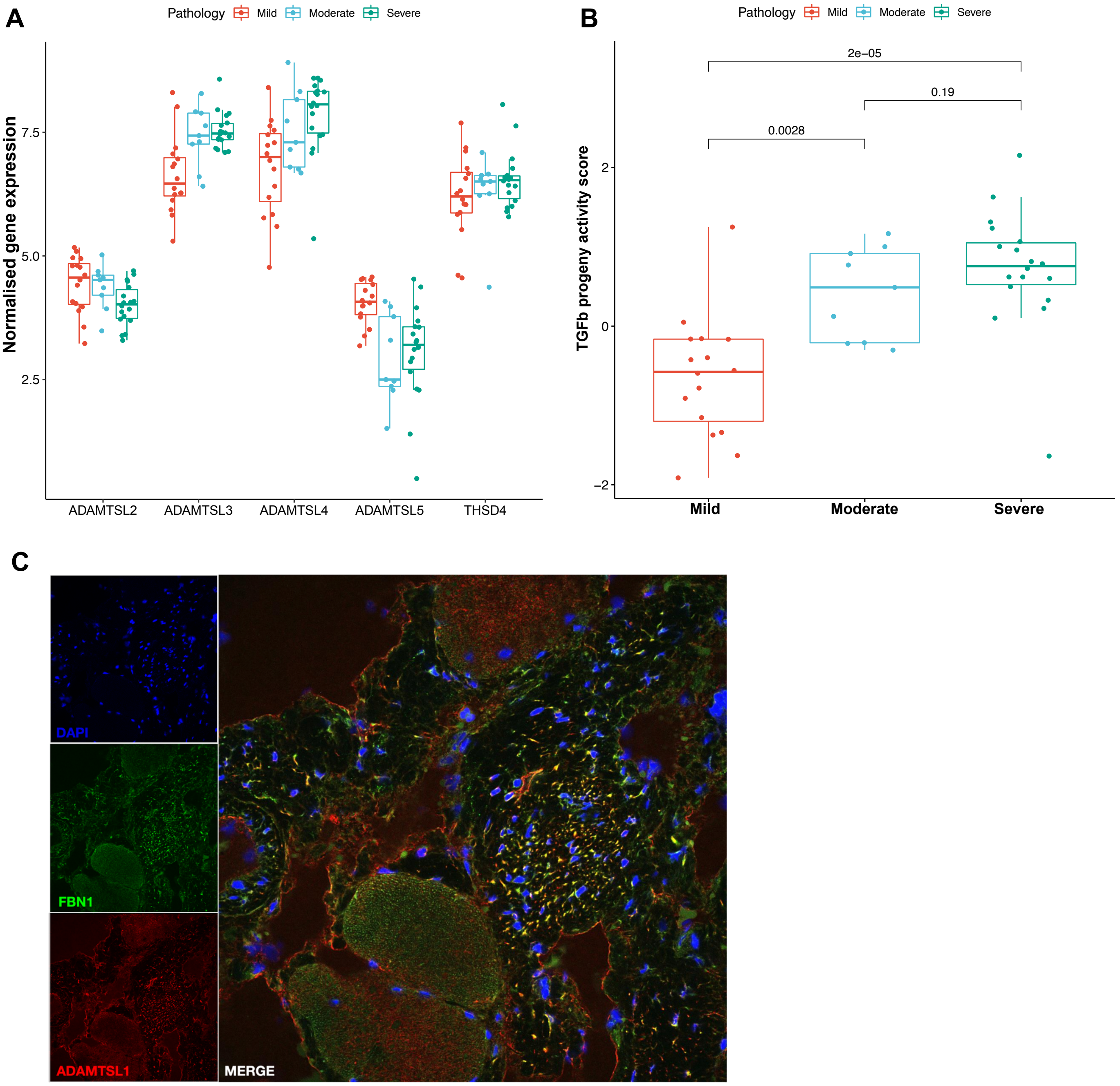

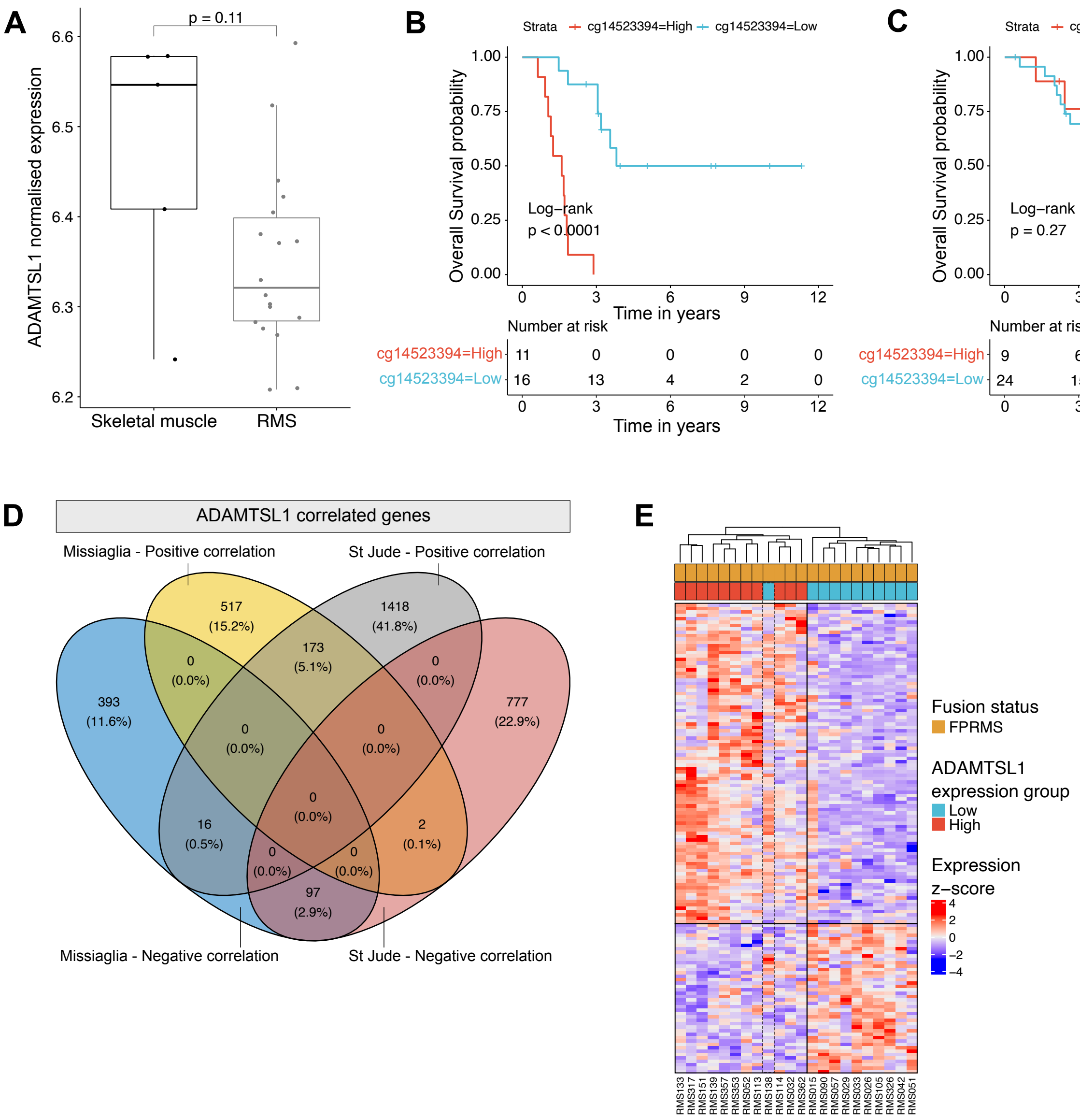
